# Large-scale genomic study reveals robust activation of the immune system following advanced Inner Engineering meditation retreat

**DOI:** 10.1101/2021.05.18.444668

**Authors:** Vijayendran Chandran, Mei-Ling Bermúdez, Mert Koka, Brindha Chandran, Dhanashri Pawale, Ramana Vishnubhotla, Suresh Alankar, Raj Maturi, Balachundhar Subramaniam, Senthilkumar Sadhasivam

## Abstract

The positive impact of meditation on human wellbeing is well documented, yet its molecular mechanisms are incompletely understood. We applied a comprehensive systems biology approach starting with whole blood gene expression profiling combined with multi-level bioinformatic analyses to characterize the co-expression, transcriptional, and protein-protein interaction networks to identify meditation-specific core network after an advanced 8-day Inner Engineering retreat program. We found the response to oxidative stress, detoxification, and cell cycle regulation pathways were downregulated after meditation. Strikingly, 220 genes directly associated with immune response, including 68 genes related to interferon (IFN) signaling were upregulated, with no significant expression changes in the inflammatory genes. This robust meditation-specific immune response network is significantly dysregulated in multiple sclerosis and severe COVID-19 patients. The work provides a foundation for understanding the effect of meditation and potential implications to voluntarily and non-pharmacologically improve the immune response before immunotherapy for many conditions, including multiple sclerosis and COVID-19 vaccination.

## Introduction

Yoga and meditation are holistic disciplines that integrate both mental and physical methods for human wellbeing [1]. These practices are growing in popularity worldwide, and according to a recent national health survey, 14% of the adult United States population used yoga or meditation within the previous year [2]. Several studies have demonstrated multiple health benefits from such methods [3, 4], including reduced stress [5-8], anxiety [5, 7, 9, 10], fatigue [5, 11], depression [5, 9, 12], chronic pain [13-15], and disease severity for inflammatory bowel disease [16, 17] and cardiovascular disease [6, 18, 19]. However, the mechanisms responsible for these improvements are poorly understood. These parameters are typically measured with self-reported surveys before and after meditation interventions, and such an approach may be prone to bias and subjectivity. Several studies on meditative practices have, however, shown changes in gene expression levels demonstrating that these methods may benefit physiology at its most fundamental level [20, 21]. Nevertheless, most of the previous studies are: (a) cross-sectional studies (evaluating only one time-point) [22-26], (b) done on highly experienced meditators [22, 23], (c) small sample-sized [22-24, 26, 27], (d) tested on handpicked non-specific biomarkers [25, 26], and (e) confounded with different lifestyle and diet [22, 25, 26].

To understand the meditative effect and to overcome these limitations, (a) we applied unbiased gene expression analyses on 4 time-points before and after the intensive 8-day Samyama meditation (an advanced Inner Engineering program attended by approximately 20,000 participants to date), and (b) analyzed the transcriptomic changes from 388 samples obtained from 106 individuals after a residential meditation retreat including a vegan diet at The Isha Institute of Inner Sciences (McMinnville, TN). Participants spent 8 days in complete silence with more than 10 hours of meditation per day. We reasoned that improvement in multiple physical and mental health conditions by this meditative practice would likely reflect significant differences in intrinsic transcriptional networks, rather than changes in expression of a few individual genes. We applied a multi-staged approach to characterize the co-expression, transcriptional, and protein-protein interaction networks associated with meditation, and we validated several network predictions using bioinformatic approaches.

The co-expression network that is upregulated after meditation consisted of 220 genes directly related to immune response, including 68 genes directly associated with interferon signaling. Transcription factor binding site enrichment analysis for this network also revealed six interferon regulatory factors (IRF1, 2, 3, 7, 8, and 9) binding sites were over-represented. By integrating protein-protein interactions with the RNA co-expression network, we identified a core meditation-associated network highly enriched with core set of hub genes, including well-known regulators known to promote immune function and novel regulators not previously associated with meditation. Our analysis indicates that, rather than acting in isolation, these enriched regulators provide crucial cross-talk between the activated functional pathways after meditation. We show that 24 drivers directly related to interferon alpha and gamma pathways are upregulated after meditation. Moreover, we show that the expression of 3 transcription factors (TFs; *STAT1, STAT2*, and *TRIM22*) is coordinately upregulated following meditation and regulating the majority of genes enriched for gene sets related to interferon signaling. This is consistent with the notion that coordinate regulation of this core network, rather than individual components, is necessary for interferon signaling activation after meditation. We also show that these 24 interferon-related drivers and their targets are also upregulated in COVID-19 patients with mild disease, but significantly downregulated in cases with severe disease. We observed the opposite trend for cytokine and inflammation signaling genes, as we found significant upregulation in severe compared with mild COVID-19 patients (and no significant changes after meditation). Finally, by comparing the transcriptional changes in multiple sclerosis (MS) patient samples, we showed that these critical meditation-specific drivers regulating the interferon signaling were dysregulated in untreated MS patients and eventually elevated in MS patients after IFN disease-modifying treatment. This study highlights meditation as a behavioral intervention that could have important implications for treating various conditions associated with excessive or persistent inflammation with a dampened immune system profile.

## Results

### Blood-based genomic analysis reveals large scale genetic changes linked to intense meditation

The Samyama retreat was conducted in April 2018 at the Isha Institute of Inner Sciences (McMinnville, TN) with the requirement to remain entirely silent for 8 days, with more than 10 hours of meditation per day, a strict vegan diet, and a regular sleep-wake cycle. The study exclusion criteria included inability to read or comprehend the consent form; subjects with severe anemia; active use of marijuana, opioids, or related drugs; use of antibiotics or probiotic/prebiotic supplements within 60 days of enrollment; and participants living outside of the country. Blood specimens (196 from female and 192 from male) were collected at 4 different times from participants with an average of 40 years of age for both sexes (Fig. 1A). In total, we characterized the transcriptome profiles of whole-blood cells from 388 specimens obtained from 106 individuals before and after the meditation retreat at 4 time-points (T1–T4) by RNA sequencing (RNA-Seq). The T1 samples were collected 5-8 weeks before the retreat, T2 samples were collected on the day of retreat before starting the meditation method, T3 samples were collected immediately after the retreat, and T4 samples were collected three months after the retreat (T1= 97, T2= 105, T3= 104, T4= 82; N= 388) (Fig. 1A). First, we performed cellular deconvolution to identify cell types and examine their heterogeneity in composition between the time points [28]. Interestingly, out of 22 different blood cell types, we observed higher neutrophil and lower CD4 and CD8 T cell relative proportions at T3 than other time points (Fig. S1).

**Fig. 1.**
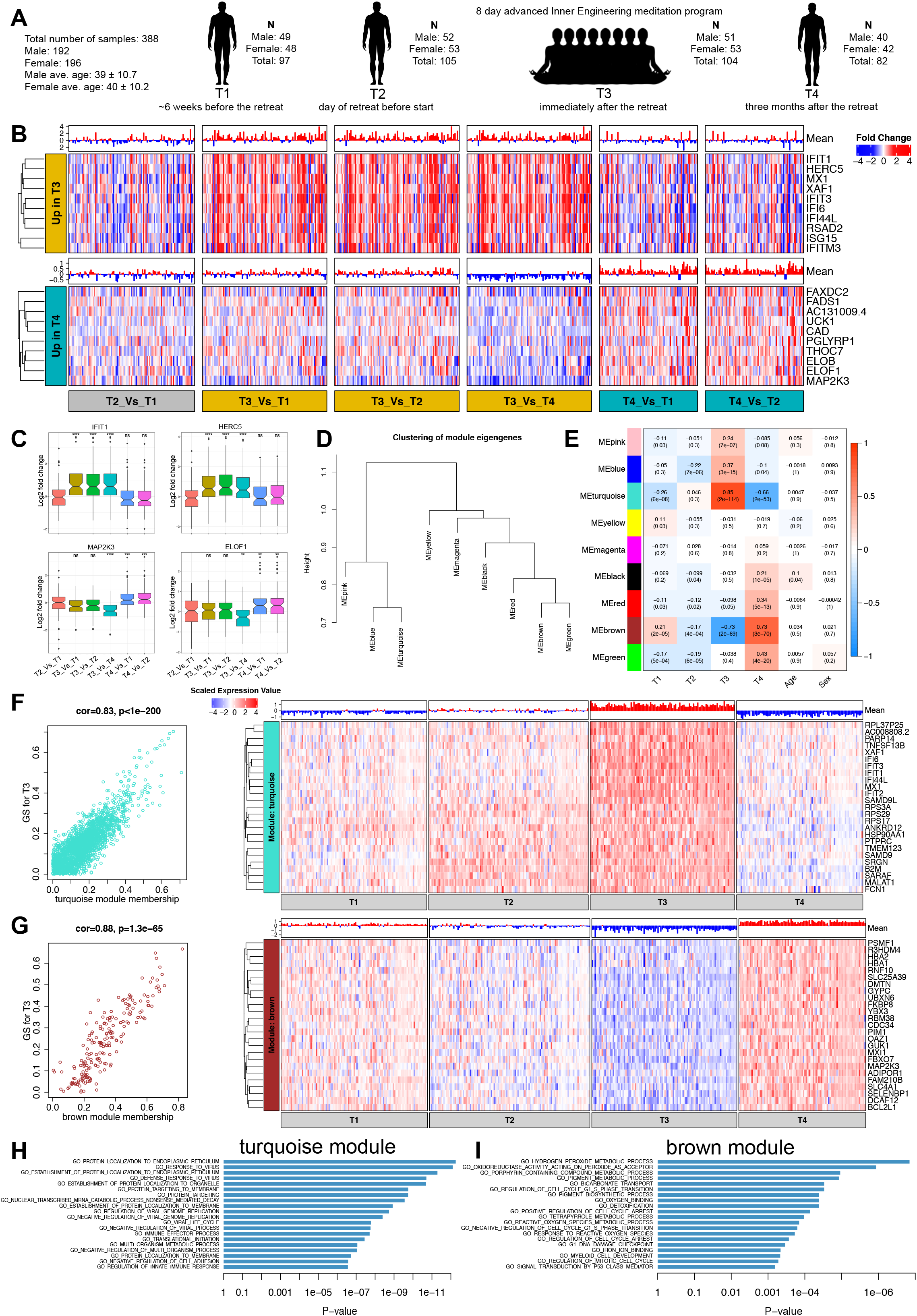
Co-expression network analysis of whole blood transcriptome profile changes after meditation. (**A**) Longitudinal blood collection at four time points (T1-T4) from advanced Inner Engineering meditation program participants. (**B**) Heatmaps are depicting the expression of the top ten genes (rows) across samples (columns) for six different timepoint comparisons (red corresponds to gene upregulation and blue to downregulation). Mean gene expression levels are shown as a bar-plot. (Top and bottom panel) The top 10 genes upregulated at T3 (after meditation) and T4 (3 months after meditation) are shown. (**C**) Boxplot represents the variability in the expression levels of the top two genes upregulated at T3 (top) and T4 (bottom). (**D**) Hierarchical clustering of WGCNA module eigengenes. (**E**) Module-trait associations. Each row corresponds to a module eigengene, column to a trait. Each cell contains the corresponding correlation and p-value. Red and blue color note positive and negative correlation with gene expression, respectively. (**F, G**) A scatter plot of gene significance (GS) for T3 versus the module membership (MM) in the turquoise (top) and brown (bottom) module. Intramodular analysis of the genes found in these two modules contains genes that have a high correlation with meditation, with p<1.3e-65 and correlation >0.8. Heatmaps are depicting the expression of top 25 hub genes (rows) across samples (columns) for four timepoints (red corresponds to gene upregulation and blue to downregulation) in the turquoise (top) and brown (bottom) module. Mean gene expression levels are shown as a bar-plot. Hub genes have a high GS and MM value with T3. (**H, I**) Gene set enrichment analysis of Gene Ontology terms calculated by Fisher’s exact test for the turquoise and brown module.

To study transcriptome changes linked to blood cells, we eliminated any confounding effects associated with this cell-type heterogeneity. We used the cell type proportion estimates to adjust the data for all analyses (see Methods). Next, we performed differential expression analyses on batch and cell type composition corrected data in all 4 timepoints (T1, T2, T3, T4) (see Methods) by pair-wise comparison of all 6 permutation combinations and identified 1649 genes differentially expressed in total (Table S1). We observed distinct early (T3) versus late (T4) genetic alterations in blood cells ‘after’ meditation. Out of 1649 differentially expressed (DE) genes, we observed almost 44% (719) of DE genes exclusively in the timepoint immediately after meditation (T3), followed by 30% (496) differentially expressed at the 3-month follow-up (T4) compared with pre-meditation timepoints (T1 or T2). We observed 116 genes differentially expressed only in T3 (Table S1), and the top 10 DE genes (*IFIT1, HERC5, MX1, XAF1, IFIT3, IFI6, IFI44L, RSAD2, ISG15, IFITM3*) in T3 alone were directly related to immune response including antiviral defense function (Fig. 1B, C). This suggests a previously unknown influence of meditation intervention on the genetic make-up of blood cells with the early wave associated immune-related changes (Table S1 and Fig. 1B). We also observed long-term effects due to meditation; 58 genes were differentially expressed only in T4 (3 months after meditation), which consisted of several upregulated genes involved in catalytic activity (*CAD, FADS1, FAXDC2, MAP2K3, PGLYRP1*, and *UCK1*), localized to the mitochondrial membrane (*CISD2, NDUFAF1*, and *MRPS18A*), and involved in translation elongation factor activity (*ELOF1* and *ELOB*) (Fig. 1B, C). Together, these findings suggest that meditation has an immediate impact on immune cells and genes, which are transient in nature and followed by a dynamically altered effect observed 3 months after the meditation retreat.

### Gene co-expression network analysis identify modules associated with meditation

We next performed weighted gene co-expression network analysis (WGCNA) [29, 30], a powerful method for understanding the modular network structure of the transcriptome, to organize the meditation related transcriptional changes in an unbiased manner. WGCNA permits the identification of modules of highly co-expressed genes whose grouping reflects shared biological functions and key functional pathways and key hub genes within the modules, and has been widely applied to understanding disease-related transcriptomes [31, 32]. Using WGCNA [30, 32], we identified 9 robust co-expression modules (Fig. 1D, E, and Table S2) for the datasets generated at 4 time-points before and after meditation. Based on the significant module trait (time-points) relationships, we identified 2 modules (turquoise and brown) strongly associated with T3 (Fig. 1D, E). To further validate turquoise and brown modules, we calculated gene significance (correlation of a gene expression profile with a sample trait) with T3 and observed significant correlation (Fig. 1F, G). Based on the eigengene-based intramodular connectivity (module membership), we observed that genes present in the turquoise module were significantly upregulated and genes present in the brown module were significantly downregulated following meditation (Fig. 1H, I). These 2 modules with a high correlation between gene significance and module membership could represent pathways associated with meditation. Gene ontology (GO) enrichment analysis revealed that several categories related to immune function, protein targeting, and localization were upregulated at T3. The upregulated RNA co-expression network (turquoise module) included many previously identified genes known to regulate the immune system and related pathways, including 220 genes differentially expressed after meditation, which are directly related to the immune response (Table S3). This list included 68 genes related to interferon signaling. Interestingly, the top 10 hub genes in the upregulated module included well-known genes directly associated with type I interferon signaling pathway, namely *IFI6, IFIT1, IFIT2, IFIT3, IFI44L, MX1*, and *XAF1* (Fig. 1F). The downregulated co-expression module was enriched for hydrogen peroxide, detoxification, and cell cycle process, including Glutathione Peroxidase 1 (*GPX1*) and Peroxiredoxin 2 and 5 (*PRDX2* and *PRDX5*) (Table S3). *GPX1* catalyzes the reduction of hydrogen peroxide by glutathione and thereby protects cells against oxidative damage. *PRDX2* and *PRDX5* play a role in cell protection against oxidative stress by detoxifying intracellular peroxides. These results suggest that meditation leads to differential expression of several genes related to oxidative stress, in agreement with a previous study [33]. The downregulated module also included genes regulating the cell cycle, namely, BCL2 Like 1 (*BCL2L1*), F-box protein 7 (*FBXO7*), Pim-1 proto-oncogene serine/threonine kinase (*PIM1*), and RNA binding motif protein 38 (*RBM38*). BCL2L1, is known to contribute to programmed cell death [34], PIM1 is known to control cell growth, differentiation, and apoptosis [35], RBM38 is shown to regulate cell-cycle arrest [36], and FBXO7 is shown to activate cell cycle regulators [37]. Downregulation of these genes suggests that meditation may exert effects on cell cycle regulation through transcriptional regulation.

### Transcriptional regulatory network analyses identify critical drivers associated with meditation

To identify critical regulators related to meditation, we performed NetBID analysis, a data-driven network-based Bayesian inference that identifies key drivers (TFs or signaling factors) in a given transcriptome [38]. First, we reverse-engineered a meditation-specific regulatory network using the SJARACNe algorithm [39]. We then applied the activity inference algorithm in NetBID to identify drivers whose network activity is significantly different before versus after meditation. We identified 90 drivers with significant differential activity (DA) compared to T3 versus other time-points (Fig. 2A and Table S4). To interpret the significance of these drivers we evaluated the expression pattern of their targets by plotting them on a ranked gene list (Fig. 2B). We observed that positively-regulated target genes of the upregulated DA drivers tended to have higher fold-change expression values in T3 compared with other time points. We observed the opposite pattern for the targets regulated by the downregulated DA drivers, validating the significance of these drivers and their targets (Fig. 2B-D). Next, we performed gene set enrichment analysis (GSEA) against the collection of annotated gene sets from MSigDB [40] to elucidate the functional relevance of these drivers. Strikingly, we observed several functional categories associated with immune function significantly enriched for upregulated drivers (Fig. 2E, Table S5). Among the top enriched functional categories included interferon-alpha and gamma response and other direct immune-related categories (Fig. 2E). Next, we examined the enriched functional categories associated with the target genes of these drivers utilizing the hallmark gene sets from MSigDB. Remarkably, we observed that the targets of these drivers were significant enriched in interferon-alpha and gamma response genes sets (Fig. 2F). We also observed 24 of these upregulated DA drivers were also present within the hallmark interferon alpha and gamma response genes sets from MSigDB (Fig. 2G). These 24 drivers included interferon-induced protein with tetratricopeptide repeats (*IFIT1, IFIT2*, and *IFIT3*), interferon stimulated gene 15 (*ISG15*), interferon-induced MX Dynamin Like GTPase 1 (*MX1*) with known antiviral activity. *MX1* gene protects mice against influenza strains pandemic 1918 and the highly lethal human H5N1 [41]. Other drivers included interferon-induced production of the 2’-5’-oligoadenylate synthetase family (*OAS1, OAS2*, and *OAS3*) and DExD/H-Box Helicase 58 and 60 (*DDX58* and *DDX60*). RNA helicases DDX58 are known to activate kinases which phosphorylate the interferon regulatory factor IRF7 to induce IFN-alpha and IFN-beta interferons [42, 43]. Similarly, *DDX60* is known to positively regulate *IFIH1*-dependent type I interferon signaling [44]. Together, these data suggest that immune function pathways (particularly interferon-related) are enhanced immediately after meditation.

**Fig. 2.**
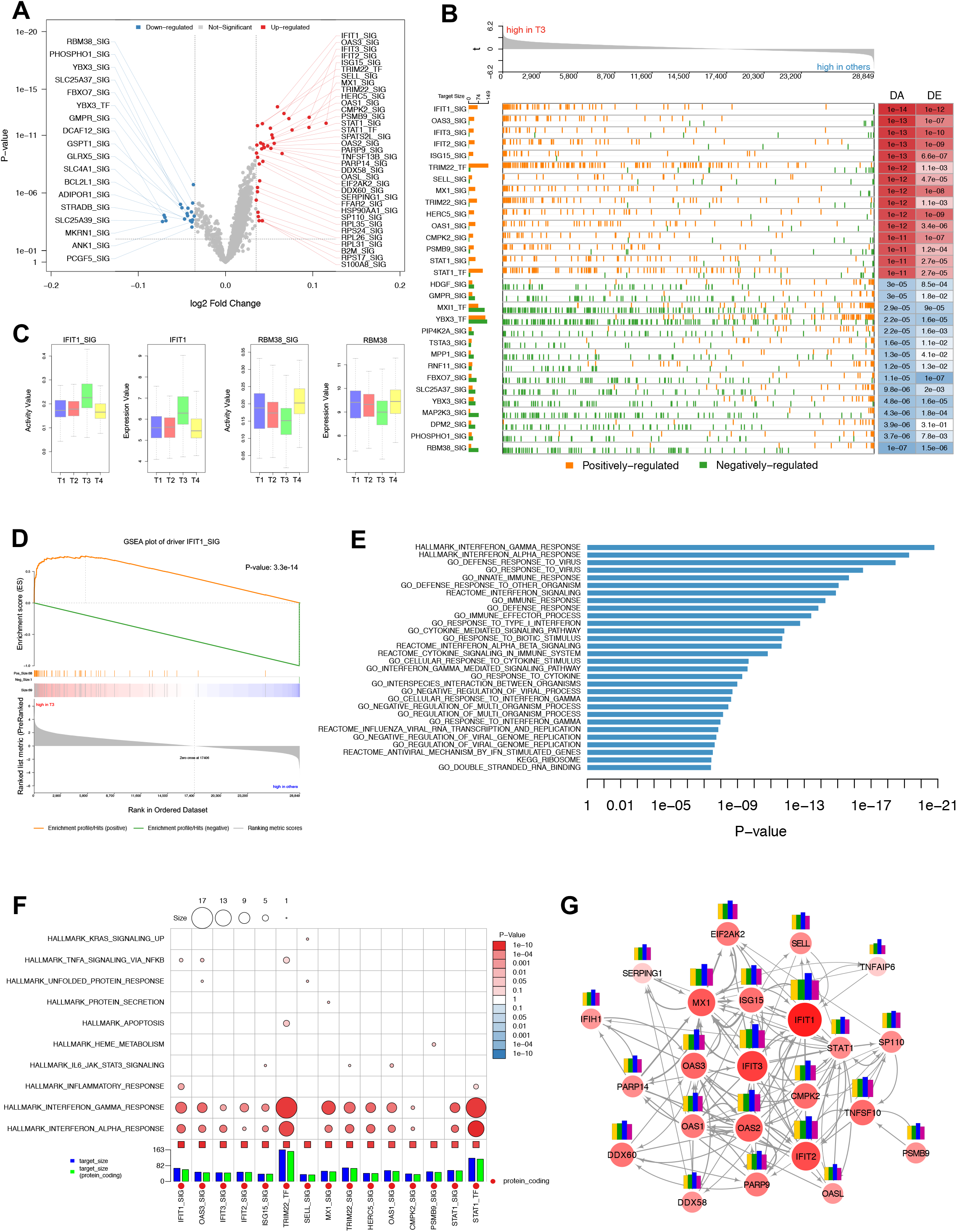
Transcriptional regulatory network analyses identify critical signaling pathways and drivers associated with meditation. (**A**) Volcano plot displaying top drivers (transcription factors (TF) or signaling factors (SIG)) with significant differential activity (DA) in T3 when compared to other time points. The red dots represent the upregulated drivers; the green dots represent the downregulated drivers after meditation. (**B**) GSEA plot shows the significance of top DA drivers with their expression and regulation of their target genes. Positively-regulated target genes (orange) of the upregulated DA drivers (red) tend to have higher fold change expression values in T3 than other time points. The opposite pattern is observed for the targets regulated by the down-regulated DA drivers (blue). The number of target genes for each driver (left) and DA and DE (differential expression) values (right) are shown. (**C**) Boxplot representation of the variability in the activity (left) and expression (right) values of top driver upregulated (*IFIT1*) and downregulated (*RBM38*) after meditation. (**D**) GSEA enrichment plot of top driver *IFIT1* shows significant enrichment for positively-regulated target genes with higher fold change expression values in T3 than other time points. (**E**) Biological function enrichment of top DA drivers. Bar charts show significantly enriched gene sets in the gene MSigDB database for the top drivers. (**F**) Biological function enrichment of the target genes of top 15 DA drivers against curated hallmark gene sets from MSigDB. (**G**) Subnetwork of the reverse-engineered meditation-specific regulatory network showing upregulated DA drivers directly associated with interferon-alpha and gamma response. Nodes correspond to genes and edges to mutual information (MI). Edge size corresponds to MI value. Larger nodes correspond to differential expression (Z-score). Node color represents upregulation (red) or downregulation (blue). Bar plot depicting the gene expression levels at different time points (left to right; T1 to T4), showing consistent elevation at T3 for all drivers.

### Transcription factor binding site enrichment in meditation specific co-expression modules

From network analyses, we observed 220 genes directly associated with immune response in the turquoise module - the major upregulated module after meditation, which contained 68 genes related to interferon signaling including the 24 IFN driver genes (Fig. 2G). To uncover the potential regulatory network contributing to the consistent co-expression of multiple immune-related genes after meditation, we performed sequence-based TF-binding site (TFBS) enrichment analysis for both turquoise and brown co-expression modules (see Methods). To avoid confounders and identify the most statistically robust sites, we used 3 different background datasets (1,000-bp sequences upstream of all human genes, human CpG islands, and the human chromosome 20 sequence). We identified 51 TFs (see Methods) significantly enriched in these 2 co-expression modules (Fig. 3A-B, Table S6). Remarkably, 6 IRF genes were over-represented in the turquoise module, the major upregulated module after meditation (Fig. 3A). Next, we examined each TF’s association with IFN signaling from published literature by testing association with the keywords “interferon signaling” or “interferon pathway” in the PubMed database for every TF (see Methods). This analysis identified 16 TFs (31%) that were strongly associated with IFN signaling, which included Signal transducer and activator of transcription 1 and 2 (*STAT1, STAT2*), and several IFN genes: *IRF1, 2, 3, 7, 8*, and *9* (Fig. 3A, Table S7). Notably, 3 of these over-represented TFs (STAT1, STAT2, and IRF7) are present in the turquoise module, the major upregulated module after meditation (Fig. 3A-B, Table S6), suggesting their critical role in activating the enriched pathways in the turquoise module. Interaction of IRFs with STAT1-STAT2 heterodimers or with STAT2 homodimers in response to IFNs redirects these complexes to a distinct group of target genes harboring the interferon-stimulated response element (ISRE) to regulate their expression levels involved in IFN signaling [45]. Together these results suggest critical involvement of these TFs in IFN signaling after meditation.

**Fig. 3.**
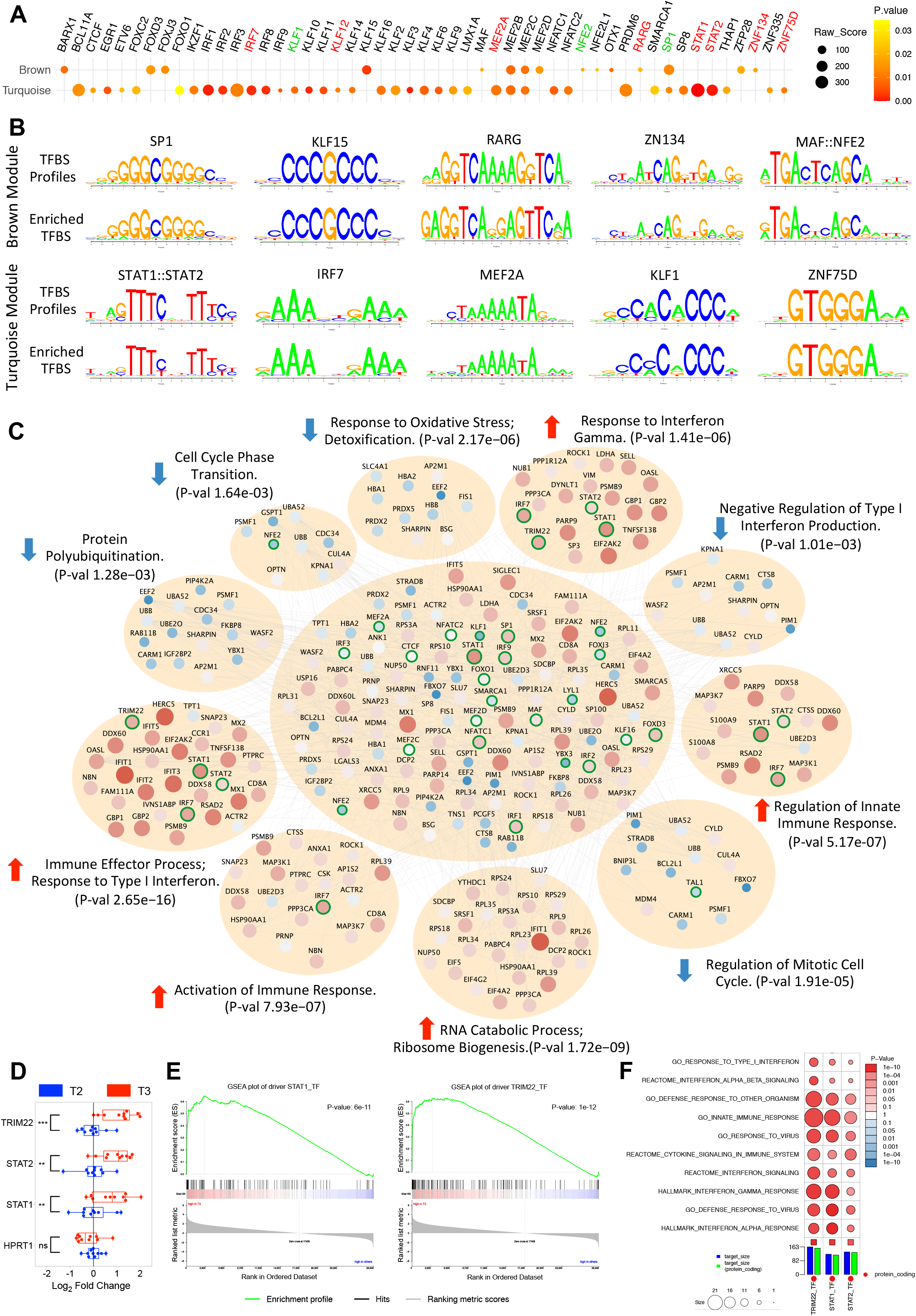
Transcription factor binding site enrichment and convergent pathway analyses. (**A**) Dot plot showing 51 enriched TFs for the brown (top) and turquoise (bottom) module. The dot’s size corresponds to the raw enrichment score obtained from the Clover algorithm, and the color of the dot corresponds to enrichment significance. Highlighted names of the overrepresented TFs denote their presence in the brown (green) or turquoise (red) module. (**B**) Sequence logo plots of reference TFBS profiles (JASPAR/HOCOMOCO) and identified position weight matrix for selected TFs significantly over-represented and present in the brown (top) or turquoise (bottom) module. (**C**) Protein-protein interaction network represented by the genes from both turquoise and brown modules along with over-represented TFs and all drivers with significant differential activity. Significantly enriched functional pathways (corrected P-value < 0.05 from Fisher’s exact test) in the PPI network are shown. Nodes correspond to genes and edges to PPI. Larger nodes correspond to differential expression (Z-score). Node color represents upregulation (red) or downregulation (blue). Nodes with highlighted border (green) correspond to enriched or driver TFs. (**D**) Quantitative PCR validation of three critical upregulated TFs after meditation. Boxplot representation of the expression levels of three critical TFs from ten random individuals for each time point. Expression levels of HPRT1 is shown as endogenous control. Paired t-test was performed to calculate the statistical significance (**p ≤ 0.01, ***p ≤ 0.001). (**E**) GSEA enrichment plot of *STAT1* and *TRIM22* showing significant enrichment for positively-regulated target genes with higher fold change expression values in T3 compared to other time points. (**F**) Biological function enrichment of the target genes of *STAT1, STAT2*, and *TRIM22* against curated gene sets from MSigDB showing enrichment for several gene sets related to immune and interferon pathways.

### Co-regulated genes after meditation represent convergent pathways

To extend this work to the level of specific proteins and identify potential conserved protein-functional pathways represented by the meditation co-expression modules, we determined the PPI (protein-protein interaction) network represented by the genes in both turquoise and brown modules (see Methods). We reasoned that this would not only provide independent validation of the relationships inferred by RNA co-expression and regulatory networks but that the PPIs would allow us to dissect significant functional pathways impacted by meditation. We screened experimentally validated PPIs among all possible combinations of gene pairs present in the co-expressed modules, over-represented TFs (Fig. 3A), and all drivers with significant differential activity (Fig. 2A), obtaining a PPI network consisting of 143 unique nodes and 495 edges (Fig. 3C). We observed enrichment of critical functional pathways related to the response to oxidative stress, detoxification, regulation of cell cycle, protein polyubiquitination, and negative regulation of type I interferon production to be downregulated after meditation (Fig. 3C). Likewise, we also observed enrichment of several critical functional pathways that contribute to the immune function to be upregulated after meditation, which included immune effector process, response to type I interferon, RNA catabolic process, activation of immune response, regulation of innate immune response, and response to IFN-gamma (Fig. 3C). These results validate the network-based approaches as immune-related pathways were consistently enriched in co-expression, regulatory, and PPI network analyses. By integrating these 3 different network analyses, we found 3 critical TFs; STAT1, STAT2, and TRIM22 (tripartite motif containing 22), that are differentially expressed after meditation (Fig. 3D) and regulate several genes that are enriched for gene sets related to IFN signaling (Fig. 3E, F). All 3 TFs are known to be IFN-induced and play a role in restricting infection from diverse viruses by activating immune function [46] [47]. Together, these results suggest that meditation as a behavioral intervention could have important implications for targeting diseases with impaired IFN response.

### Comparison of transcriptional profiles obtained from COVID-19 and multiple sclerosis patient samples

Impaired IFN response and exacerbated inflammatory response are generally observed in viral infections and are the hallmark of severe SARS-CoV-2 infection [48]. Hence, we compared the transcriptional network obtained from a publicly available dataset of leukocytes from COVID-19 patients with our meditation dataset to directly examine their expression pattern. First, we examined the expression levels of well-established viral infection marker genes (*PTX3, CXCL10, CD46*) in both datasets. As expected, we observed upregulation of these marker genes in the COVID-19 dataset but not in the meditation dataset (Fig. S2). Next, we examined the expression levels of 24 meditation-associated DA drivers (Fig. 2G), which are hallmark IFN-alpha and -gamma response genes. The comparison of T3 (after meditation) versus T2 (before meditation) revealed significant upregulation of these drivers when compared with T2 versus T1 (baseline) (Fig. 4A). The comparison of COVID-19 patients (non-intensive care unit [ICU]) versus non-COVID-19 patients (non-ICU) also revealed upregulation of several of these drivers (Fig. 4B). Strikingly, the comparison of severe COVID-19 patients (ICU) versus mild COVID-19 patients (non-ICU) revealed downregulation of these drivers (Fig. 4B). This suggests impairment of these 24 critical drivers regulating IFN signaling in severe COVID-19 patients, which are elevated after meditation. Next, we examined the expression of the targets of these 24 drivers, which are also part of the hallmark IFN-alpha and -gamma response gene sets from MSigDB. For this analysis, we merged the reverse-engineered regulatory network generated for both COVID-19 and meditation datasets to examine the changes. We observed that 97% of these genes were upregulated after meditation, 76% of these genes were upregulated in mild COVID-19 patients, and, strikingly, only 31% of IFN response genes were upregulated in severe COVID-19 patients (where the majority of the genes were downregulated) (Fig. 4C). We also found 8 upregulated DA drivers (*ARG1, CD55, HMGB2, IL18R1, IL1R1, IL1R2, IRAK3, LTF*) associated with cytokine and inflammation signaling in COVID-19 regulatory network. By examining the expression of the targets of these drivers present in the hallmark inflammatory response gene set from MSigDB, we observed significant upregulation of these inflammation-related genes in severe COVID-19 patients compared with mild COVID-19 patients (Fig. 4D). The comparison of T1 (baseline), T2 (before meditation), and T3 (after meditation) revealed no significant changes in these inflammation-related genes (Fig. S3A). Previous studies have shown that exercise also improves immune competency [49]. By comparing the whole-blood transcriptome datasets of exercise training with the meditation dataset, we did not observe upregulation of the 24 IFN driver genes after exercise training suggesting a unique gene signature associated with meditation (Fig. S3B, C). These results suggest that a unique gene signature associated with meditation enhances immune function without activating inflammatory signals impaired in severe COVID-19 patients.

**Fig. 4.**
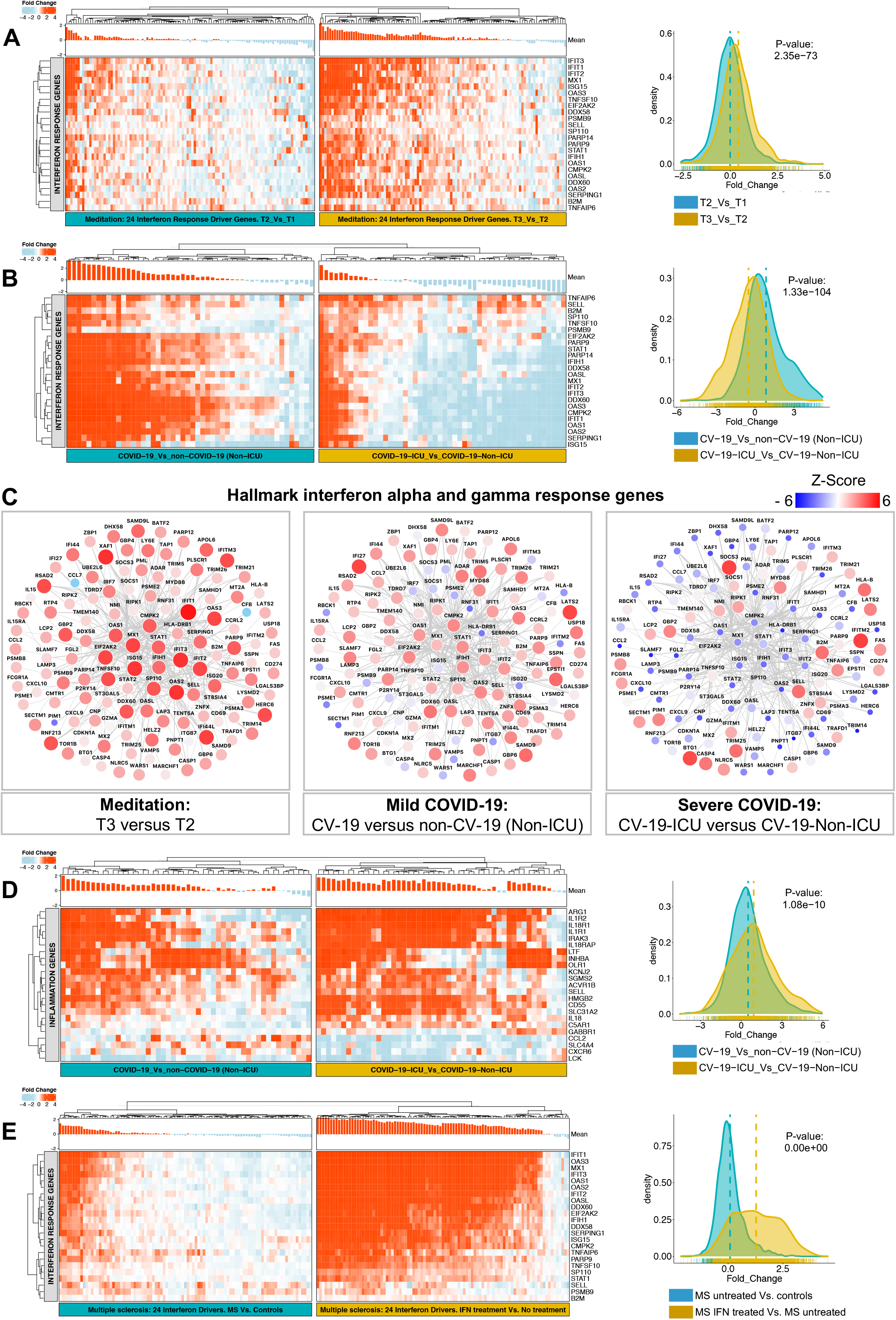
Comparison of interferon and inflammatory response in meditation, COVID-19 and multiple sclerosis specific transcriptional profiles. (**A**) Heatmaps are depicting the expression of the 24 interferon response driver genes (rows) across meditation samples (columns) for two different timepoint comparisons, T2 (before meditation) versus T1 (baseline) and T3 (after meditation) versus T2. (**B**) Heatmaps comparing the expression of the 24 IFN driver genes in COVID-19 samples for two different comparisons; non-ICU COVID-19 patients versus non-ICU non-COVID-19 patients (left – mild patients) and ICU COVID-19 patients versus non-ICU COVID-19 patients (right – severe patients). (**C**) Subnetwork of the reverse-engineered meditation-specific and COVID-19-specific regulatory network showing targets of the 24 IFN drivers directly associated with interferon-alpha and gamma response, displaying dramatic downregulation in severe COVID-19 patients. Nodes correspond to genes and edges to mutual information (MI). Larger nodes correspond to differential expression (Z-score). Node color represents upregulation (red) or downregulation (blue). (**D**) Heatmaps comparing the expression of the 23 inflammatory response genes in COVID-19 samples for two different comparisons as described in **B**, displaying significant upregulation in severe COVID-19 patients. (**E**) Heatmaps comparing the expression of the 24 meditation-specific IFN driver genes in multiple sclerosis samples for two different comparisons; MS patients versus controls (left) and MS patients after IFN treatment versus MS untreated patients (right). For all heatmaps, the red color corresponds to gene upregulation and blue to downregulation. Mean gene expression levels are shown as a bar-plot on top of each heatmap. All the heatmaps are supplemented with density plots showing the distribution of log2 fold change, and the significance of the variability in the expression levels between the two groups are calculated by a two-sample t-test.

Next, we compared the transcriptional changes in a publicly available multiple sclerosis (MS) dataset. MS is characterized by autoimmune inflammation and subsequent neuronal degeneration. IFN treatment was the first disease-modifying therapy available to treat MS, preventing and reducing relapse rates and reducing new brain lesions, progression, and cognitive loss [50]. Hence, we examined the expression levels of 24 meditation-associated IFN response driver genes in the whole-blood gene expression dataset obtained from MS patients undergoing β-IFN therapy. Firstly, we observed that the upregulated meditation-associated IFN response driver genes were significantly downregulated in MS versus control comparison (Fig. S3D). Strikingly, the comparison of MS IFN-treated versus MS untreated patients revealed dramatic upregulation of these drivers compared with MS untreated versus controls (Fig. 4E). These results revealed that upregulation of these 24 critical drivers regulating IFN signaling in MS patients after IFN disease-modifying treatment is also elevated after meditation. Together, these results support the hypothesis that meditation contributes to improving multiple health conditions by regulating various critical pathways directly related to the disease pathogenesis.

## Discussion

Yoga and meditation techniques have improved health outcomes and quality of life for patients with many disease conditions. No comprehensive study had previously been conducted to evaluate how meditation impacts biological processes that are directly involved in disease pathogenesis. We evaluated an advanced level 8-day meditation practice and applied a comprehensive systems genomics approach, including gene expression profiling combined with multi-level bioinformatic analyses and validation of network predictions. We hypothesized that meditative practices would likely impact intrinsic transcriptional networks directly related to the disease pathogenesis rather than solely change the expression of a few individual genes. Pathways related to immune system activity are particularly relevant in this context, given that immune function has been implicated in several major health conditions, including asthma, cardiovascular disease, certain types of cancer, depression, metabolic disorders, multiple sclerosis, neurodegenerative disorders, obesity, osteoporosis, posttraumatic stress disorder, rheumatoid arthritis, and sepsis [51, 52]. In this study, using an integrative approach we found several immune-related and other fundamental cellular pathways were altered after the meditation retreat. This work provides a key proof of principle for the power of the systems genomic approaches in tackling complex problems by demonstrating that meditation practices enhance gene networks associated with distinct pathways.

In this study, we uncovered a novel immunomodulatory function of meditation via the activation of IFN pathways and observed higher neutrophil relative proportions after meditation, which accompanied the elevated interferon pathway activity. Our gene co-expression network analysis identified upregulated and downregulated modules with a significant correlation with meditation. The upregulated RNA co-expression network included many previously identified genes known to regulate the immune system and related pathways, including 220 genes differentially expressed after meditation, directly related to the immune response. This list included 68 genes related to IFN signaling. The IFN signaling pathway plays a critical role in immune function by triggering a complex regulatory system of innate and adaptive immune responses to defend against pathogens and diseases. Likely, countering a dysregulated immune system profile with meditation could theoretically function to improve health outcomes by enhancing immune defenses that protect against viral and bacterial infection and various diseases with diminished immune function. The downregulated co-expression module showed enrichment for hydrogen peroxide, detoxification, and cell cycle process. These results suggest meditation differentially express several genes related to oxidative stress in agreement with a previous study [33]. The downregulated module also included genes regulating the cell cycle, suggesting that meditation may exert effects on cell cycle regulation through transcriptional regulation, in agreement with previous studies [20].

We also observed long-term effects due to meditation; 58 genes were differentially expressed only in T4 (3 months after meditation), consisting of several upregulated genes involved in catalytic activity, localized to the mitochondrial membrane, and involved in translation elongation factor. These results indicate meditation may exert long-term changes in the transcriptional profiles related to the most fundamental cellular pathways. Interestingly, most of the immune-related upregulated genes at T3 were reversed at T4, suggesting acute but not chronic immune activation due to meditation. Chronic immune activation is known to cause immune hyporesponsiveness, anergy [53], and malignancy [54]. We identified 90 drivers with significant differential activity, which regulates this robust acute immune activation after meditation. These drivers comprised 24 upregulated DA drivers associated with IFN-alpha and -gamma response signaling, including *IFIT1*-3, *ISG15, MX1, OAS1-3, DDX58, and 60. MX1* gene has been shown to protect mice against the pandemic 1918 and highly lethal human H5N1 influenza viruses [41]. DDX58 and DDX60 are known to regulate IRF7 and IFIH1 to induce IFN-alpha and -beta signaling [42-44]. Remarkably, *IRF7* and *IFIH1* are also upregulated after meditation, and TFBS enrichment analyses also revealed an over-representation of 16 TFs strongly associated with IFN signaling, which included IRF7, suggesting an active role of *IRF7* and *IFIH1* in IFN signaling after meditation. Other TFs included STAT1, STAT2, and several IFN regulatory factor genes: IRF1, 2, 3, 7, 8, and 9. Interestingly, *STAT1* and *STAT2* are present in the turquoise module, the major upregulated module after meditation, suggesting critical involvement of these TFs in IFN signaling after meditation.

By integrating protein-protein interactions with the identified networks, we characterized a core meditation-specific network highly enriched in critical pathways. The response to oxidative stress, detoxification, regulation of cell cycle, protein polyubiquitination, and negative regulation of type I IFN production were downregulated after meditation. Likewise, the activation of immune response, regulation of innate immune response, and response to IFN pathways were upregulated after meditation. From this core network analysis, we also found 3 TFs; *STAT1, STAT2*, and *TRIM22*, which are differentially expressed after meditation, regulate several genes enriched for gene sets related to IFN signaling. It is well known that by interacting with their specific receptors, IFNs activate STAT complexes. This STAT-containing complex binds to an IFN-stimulated response element in IFN-stimulated gene promoters to driver type I IFN-stimulated transcriptional activation, which is an essential primary barrier for virus infection and for activation of innate and adaptive immune responses [46]. Both *STAT1* and *STAT2* deficient mice were highly sensitive to diverse viruses and bacterial pathogens, including vesicular stomatitis virus, influenza virus, herpes simplex virus, dengue virus, and others, highlighting the importance of these TFs in antiviral immunity [46]. Likewise, we found that the IFN-induced tripartite motif protein TRIM22 is known to play a role in the restriction of infection by diverse viruses, including HIV, encephalomyocarditis virus, influenza A virus, hepatitis B virus, and hepatitis C virus [47]. Together, upregulation of *STAT1, STAT2*, and *TRIM22* by meditation may function as a host anti-viral factor induced by interferons to restrict infection by diverse viruses. Together, these results support the hypothesis that meditation contributes to improving multiple health conditions by regulating various critical pathways directly related to the disease pathogenesis.

We also showed that the meditative practice enhanced immune function without activating inflammatory signals. This suggests that meditation, as a behavioral intervention, may be an effective component in treating diseases characterized by increased inflammatory responsiveness with a weakened immune system. It has been shown that dysregulation of interferon response coupled with exuberant inflammatory cytokine production is the defining and driving feature of COVID-19 pathogenesis [48]. It has been widely suggested that a preventive measure by the administration of IFNs to elicit a pre-existing antiviral state may block viral infection at the very early stage. As of May 5, 2021, more than 40 clinical studies of various formulations of interferon treatment plans have been registered on clinicaltrials.gov for COVID-19. Interesting, early IFN treatment before peak viral replication protects mice from lethal severe acute respiratory syndrome coronaviruses (SARS-CoV) or the Middle East respiratory syndrome-coronavirus (MERS-CoV) challenge, whereas late IFN treatment failed to effectively inhibit virus replication and aggravated immunopathology [55, 56]. Daily IFN-alpha nasal drops along with standard personal protective equipment were shown to protect at-risk health-care workers from COVID-19 over 28 days [57]. These results demonstrate the benefits of early intervention or activation of the IFN response can prevent susceptible healthy people from acquiring COVID-19. By comparing the reverse-engineered meditation-specific and COVID-19-specific regulatory network, we showed that the meditation-associated upregulated interferon-related drivers and their targets were significantly downregulated in severe COVID-19 patients. We also observed the opposite trend for cytokine and inflammation signaling genes, where we observed significant upregulation in severe COVID-19 patients compared with mild patients and no significant changes after meditation. In MS patients, IFN treatment was the first disease-modifying therapy available, reducing relapse rates and disease progression [50]. By comparing the transcriptional changes in MS patient samples, we show these critical meditation-specific drivers regulating IFN signaling were also elevated in MS patients after IFN disease-modifying treatment (but not in untreated MS patients). We also observed that meditation did not elevate bona fide viral infection marker genes and displayed distinct gene signature activation compared with the gene profiles after exercise training, which is known to improve immune competency [49]. Taken together, we speculate that early and short-term non-pharmacological intervention by meditation may voluntarily provoke and enhance the immune response before immunotherapy for many conditions, including MS and COVID-19 vaccination. Future studies will be needed in healthy and diseased subjects to test these scenarios to examine the associations between meditation, immune function, and disease symptomatology.

In conclusion, we identified and characterized the transcriptional program associated with advanced yoga and meditation practices, and we bio-informatically integrated various networks to identify meditation specific core network and validate several network predictions. This core network links several immune signaling pathways via a known set of drivers, and we showed that this core transcriptional profile is dysfunctional in multiple sclerosis and severe COVID-19 infection. Very importantly, we demonstrated that these yoga and meditative practices enhanced immune function without activating inflammatory signals. This present proof-of-principle study demonstrated that the immune system can be voluntarily influenced by non-pharmaceutical interventions like yoga and meditation. This suggests that meditation as a behavioral intervention could have important implications for treating various conditions associated with excessive or persistent inflammation with a dampened immune system profile.

## Materials and methods

### Subject recruitment

The Isha Institute of Inner Sciences (McMinnville, TN) provided a list of registrants for the April 2018 Samyama Program (8-day advanced Inner Engineering meditation retreat). Invitation letters with study information, including a link to the online survey, were sent electronically to all registrants 2-3 months prior to the program. Study eligibility criteria included: advanced meditation program participants of at least 18 years of age. Study exclusion criteria were: inability to read or comprehend the consent form; subjects with medical conditions in which a blood draw would be contraindicated (e.g., severe anemia); active use of marijuana, opioids, or related drugs; use of antibiotics or probiotic/prebiotic supplements within 60 days of enrollment; participants living outside of the country. Potential subjects were provided a study information sheet at the beginning of the online survey. They were given the following options: (1) survey-only participation (no blood sampling); (2) survey and blood sampling (requiring 2 blood samples prior to the Samyama program and 2 blood samples upon completion of the program); and (3) no participation. This study was reviewed and approved by the Institutional Review Board of the Indiana University School of Medicine (IRB #1801728792). The genome-wide transcriptome analyses part of the study was also reviewed and approved by the Institutional Review Board of the University of Florida (IRB #201802368). This study was registered at ClinicalTrials.gov (NCT04366544) and complied with Consolidated Standards of Reporting Trials (CONSORT).

### Details of Samyama Program

The Samyama retreat was conducted in April 2018 at the Isha Institute of Inner Sciences (McMinnville, TN) with a strict vegan diet and sleep-wake cycle. Sixty days before the program, Samyama participants began preparatory training and dietary restrictions including vegan diet and elimination of substances such as garlic, onion, chili, eggplant, asafoetida, coffee, tea, alcohol, cigarettes, and other stimulants, intake of minimum 50% natural foods, and focus on “living” and uncooked foods. The preparatory daily practices included Hata yoga (Surya Kriya and Yogasanas; asanas, or postures, meant to knead and strengthen the body), Kriyas (Shakti Chalana Kriya and Shambhavi Mahamudra; combinations of posture, breath, and sound that are meant to purify and enhance the flow of one’s energies while simultaneously increasing general stability), Shoonya (conscious non-doing meant to bring stillness and stimulate the release of physical, mental, and emotional blocks), Pranayama (Sukha Kriya; regulation of breath, meant to facilitate overall stability, balance, and steady attention), and Ardha Siddhasana (an asana in which one sits cross legged with the heel of the left foot placed at the foundational junction of one’s energy channels, meant to help participants sit in stillness with spine erect for longer durations). During the Samyama retreat, individuals were expected to remain entirely silent for eight days, with more than 10 hours of meditation per day.

### Blood sampling and storage

Study subjects who elected blood analysis had approximately 10 mL peripheral blood taken at 4 time points (T1, T2, T3, T4). T1 samples were collected 5-8 weeks before the retreat, T2 samples were collected on the day of retreat before starting the meditation method, T3 samples were collected immediately after the retreat, and T4 samples were collected three months after the retreat. At T1 and T4, blood draws were performed at home by an in-home phlebotomist or at an Isha Center group meditation by study personnel. The remaining samples were collected at the Isha Institute of Inner Sciences (IIIS) before/after the meditation retreat. Individuals who did not submit baseline blood samples were allowed to participate at T2 and T3 (but not T4). Peripheral blood was collected in PAXgene tubes and frozen before RNA extraction. Samples were stored frozen at -80° C until analysis was performed.

### Transcriptome profiling by RNA sequencing

The human blood samples collected into PAXgene Blood RNA Tubes were removed from the -80°C freezer and incubated overnight at room temperature. A total of 389 samples from 106 individuals in 4 time-points (T1= 98, T2= 105, T3= 104, T4= 82) were included and samples were randomized before RNA extraction to eliminate any effects based on time-point, age, sex, or batch. The manufacturer’s protocol was followed for manual purification of total RNA from human whole blood (Qiagen, cat #762164). RNA was eluted with RNase free water, and the concentration and purity (A260/A280 ratios >1.8) were assessed using a spectrophotometer. RNA was also tested for suitable mass (RiboGreen) and integrity (Agilent TapeStation), reverse transcribed to complementary DNA (Lexogen QuantSeq 3′ FWD), and sequenced on a HiSeq 4000 instrument (Illumina) in the UCLA Neuroscience Genomics Core laboratory, following the manufacturers’ standard protocols. Sequencing targeted mean three million 65-nt single-stranded reads per sample, which were mapped to the human transcriptome and quantified as transcripts per million mapped reads using the STAR aligner. Briefly, all sequencing data were uploaded to Illumina’s BaseSpace in real-time for downstream analysis of quality control. Raw Illumina (fastq.gz) sequencing files were downloaded from BaseSpace, and uploaded to Bluebee’s genomics analysis platform (https://www.bluebee.com) to align reads against the human genome and to obtain raw read counts.

### Evaluation of immune cell proportions using cellular deconvolution

To analyze immune cell proportions in the whole blood samples, formatted data with gene symbols were uploaded to the CIBERSORTx web portal (https://cibersortx.stanford.edu/) and LM22 gene signature was utilized [28]. LM22 is a signature matrix file consisting of 547 genes that accurately distinguish 22 mature human hematopoietic populations isolated from peripheral blood [28]. Bulk RNA-seq data was deconvoluted using the signature matrix with bulk mode batch correction to remove variances between different platforms. Two-way ANOVA test was used to analyze differences in the abundances of different cell types before and after meditation. The p-values were corrected for multiple testing using the Benjamini-Hochberg method. P < 0.05 was considered statistically significant.

### Differential gene expression analyses

Raw data was log transformed and checked for outliers. Across samples, Pearson correlation and clustering based on variance were used as quality-control measures. One sample from T1 was not included in any analysis due to low read counts. Variance stabilizing transformation normalization method from DESeq2 package was utilized. Normalized data was processed with ‘RemoveBatchEffect’ function from ‘limma’ package for batch and cell type composition correction. Next, a linear model was fitted across the dataset, contrasts of interest were extracted, and differentially expressed genes for each contrast were selected using an empirical Bayes test statistic [58]. Differential expression analyses on cell type composition corrected or non-corrected data revealed 98% overlap in the number of genes differentially expressed. Hence, we utilized batch and cell type composition corrected data for all downstream analyses.

### Construction of co-expression networks

A weighted signed gene co-expression network was constructed using the normalized dataset to identify groups of genes (modules) associated with meditation following a previously described algorithm [59, 60]. Briefly, we first computed the Pearson correlation between each pair of selected genes yielding a similarity (correlation) matrix. Next, the adjacency matrix was calculated by raising the absolute values of the correlation matrix to a power (β) as described previously [59]. The parameter β was chosen by using the scale-free topology criterion [59], such that the resulting network connectivity distribution best approximated scale-free topology. The adjacency matrix was then used to define a measure of node dissimilarity, based on the topological overlap matrix, a biologically meaningful measure of node similarity [59]. Next, the genes were hierarchically clustered using the distance measure and modules were determined by choosing a height cutoff for the resulting dendrogram by using a dynamic tree-cutting algorithm [59]. Utilizing this network analysis, we identified modules (groups of genes) differentially expressed across different time points before and after meditation and calculated the first principal component of gene expression in each module (module eigengene). Next, we correlated the module eigengenes with time points before and after meditation to select modules for functional validation.

### Transcriptional regulatory network analyses

We applied the network-based integrative NetBID [38] algorithm to identify critical drivers associated with meditation using batch and cell type composition corrected gene expression profiles. We first reverse-engineered a meditation-specific regulatory network by using the SJARACNe algorithm [39] from 388 RNA-Seq profiles of individuals before and after meditation. We then applied the activity inference algorithm in NetBID to identify drivers whose network activity are significantly different between T3 versus other time points. In short, meditation-specific network was inferred based on the gene expression profiles based on mutual information dependency. Transcription factors and signaling factors with a high number of differentially expressed targets in the network were regarded as potential drivers. Gene set enrichment analysis (GSEA) and curated gene sets from MSigDB [40] were used to identify potential biological functions of the drivers and targets. The activity of the potential driver is calculated from the expression of its targets, and differential activities were inferred using NetBID. The drivers whose activities are significantly correlated with the timepoint T3 were utilized for enrichment analyses. We performed gene set enrichment analysis with default parameters using pathways derived from gene sets from Molecular Signatures Database [40]. All network plots were constructed using the Cytoscape software [61].

### Transcription factor binding site enrichment analyses

Transcription factor binding site (TFBS) enrichment analysis was performed using the ‘TFBSenrich’ function from ‘RegFacEnc’ package (https://tfenrichment.semel.ucla.edu/). TFBS enrichment analysis was performed by scanning the canonical promoter region (1000bp upstream of the transcription start site) for the top 200 genes (based on kME) present in the meditation-associated co-expression modules. Next, we utilized TFBS position weight matrices (PWMs) from JASPAR (746 motifs) and HOCOMOCO (769 motifs) databases [62, 63] to examine the enrichment for corresponding TFBS within each module. For TFBS enrichment all the modules were scanned with each PWMs using Clover algorithm [64]. To compute the enrichment analysis, we utilized three different background datasets (1000 bp sequences upstream of all human genes, human CpG islands and human chromosome 20 sequence). When a TFBS is over-represented (based on the P-values obtained relative to all the three corresponding background datasets) we considered it to be significant, which increases our confidence in these predictions.

### Protein-Protein Interaction (PPI) Network Analyses

We constructed an experimentally validated protein–protein interaction (PPI) network using both turquoise and brown meditation-associated co-expression gene network modules. We created all possible combinations of gene pairs present in these co-expression networks and identified all experimentally verified interaction data (in humans dataset) for their corresponding proteins in the STRING database (integration of the following databases: BIND, DIP, GRID, HPRD, IntAct, MINT, and PID) [65], constructing the protein network by force-directed layout organized by significantly enriched functional pathways (Figure 3C). Nodes correspond to genes and edges to PPI. The size of each node in the PPI network correspond to differential expression (Z-score). Node color represents upregulation (red) or downregulation (blue). Nodes with highlighted border (green) correspond to enriched or driver TFs.

### Gene Ontology, Pathway enrichment and PubMed Analyses

Gene ontology and pathway enrichment analysis was performed using the Fisher’s Exact Test with ‘funcEnrich.Fisher’ function from ‘NetBID’ package (https://github.com/jyyulab/NetBID). A list of differentially regulated transcripts for a given modules were utilized for enrichment analyses. We performed gene set enrichment analysis with default parameters using pathways derived from gene sets from Molecular Signatures Database [40]. For PubMed analyses we determined the association with the following key-words: “interferon signaling” and “interferon pathway” in the PubMed database for every gene using R (http://cran.r-project.org/).

### Quantitative real-time PCR

qRT-PCR was performed using a Bio-Rad CFX96 real-time PCR system according to the manufacturer’s instructions. Briefly, RNA was harvested from human whole blood samples and cDNA was produced using SuperScript VILO IV master mix (cat #11756050). iTaq Universal SYBR Green Supermix was used to quantify amplification of cDNA. Primers we designed using Primer-BLAST [66] and was verified to amplify one product by verifying one peak present on the dissociation curves, and standard curves were performed to show that this assay is sensitive to changes in each gene. The following primer sequences were utilized: STAT1_F: CAGCTTGACTCAAAATTCCTGGA, STAT1_R: TGAAGATTACGCTTGCTTTTCCT, STAT2_F: CCAGCTTTACTCGCACAGC, STAT2_R: AGCCTTGGAATCATCACTCCC, TRIM22_F: CTGTCCTGTGTGTCAGACCAG, TRIM22_R: TGTGGGCTCATCTTGACCTCT. Ten biological replicates (individuals) were used for each timepoints, and three technical replicates were performed for each sample. The relative expression of genes was calculated using the 2^−ΔΔCT^ method.

### Comparison of transcriptional profiles with publicly available datasets

Gene expression data sets were downloaded from the Gene Expression Omnibus (GEO), read into R, preprocessed and normalized using variance stabilizing transformation normalization method from DESeq2 package. We then calculated the correlation of gene expression between samples, and outliers with mean sample correlations more than two to three standard deviations below average were omitted until no outliers remained. Using the ‘limma’ package, a linear model was fitted across the dataset, contrasts of interest were extracted, and differentially expressed genes for each contrast were analyzed using an empirical Bayes test statistic [58]. We analyzed several publicly available different datasets and compared it with meditation dataset generated and analyzed in this study. We analyzed: (1) dataset generated from leukocyte samples from hospitalized patients with or without COVID-19 (GSE157103), dataset from whole blood of MS patients and controls (GSE41850), and exercise training datasets (GSE111554, GSE111553).

## Acknowledgements

This work was supported by the Department of Pediatrics, University of Florida. The authors appreciate support provided by Isha Institute of Inner Sciences, McMinnville, TN, for allowing to conduct this research on participants of the advanced Inner Engineering meditation retreat program. The authors appreciate support provided by the volunteers of Isha Institute of Inner Sciences for sample collection. The authors also thank thoughtful review and constructive feedback from Nagarajan Kannan, Ph.D. Department of Laboratory Medicine and Pathology, Mayo Clinic School of Medicine. The authors acknowledge Janelle S. Renschler, DVM, PhD (Department of Anesthesia, Indiana University, Indianapolis, Indiana, USA) for assistance with medical writing funded by Indiana University in accordance with Good Publication Practice (GPP3) guidelines.

## Author contributions

V.C. was involved in data generation, all data analysis and visualization, and funding acquisition. M.L.B. was involved in sample processing. M.K. was involved in sample processing and qPCR data generation. V.C., B.C., D.P., R.V., S.A., R.M., B.S., and S.S. was involved in samples collection and study design. S.S. was involved in funding acquisition for sample collection. V.C. wrote the manuscript, and all authors reviewed the manuscript.

## Data and materials availability

Datasets generated and analyzed in this study are available at Gene Expression Omnibus (GEO). Accession number: GSE174083. GEO accession number for the publicly available datasets analyzed in this study are: GSE157103 (COVID-19), GSE41850 (multiple sclerosis), and GSE111554, GSE111553 (exercise training). The WGCNA package can be found online at: https://cran.r-project.org/web/packages/WGCNA/index.html. The NetBID package can be found online at: https://github.com/jyyulab/NetBID. The TFBS enrichment package can be found online at: https://tfenrichment.semel.ucla.edu/. Any custom code generated for our analyses not specifically listed here or in the text may be requested from V.C. All R packages or other software used is given in Methods for each relevant analysis.

