## Supplementary figures and images for "Large-scale genomic study reveals robust activation of the immune system following advanced Inner Engineering meditation retreat"

### Supplemental_Figures

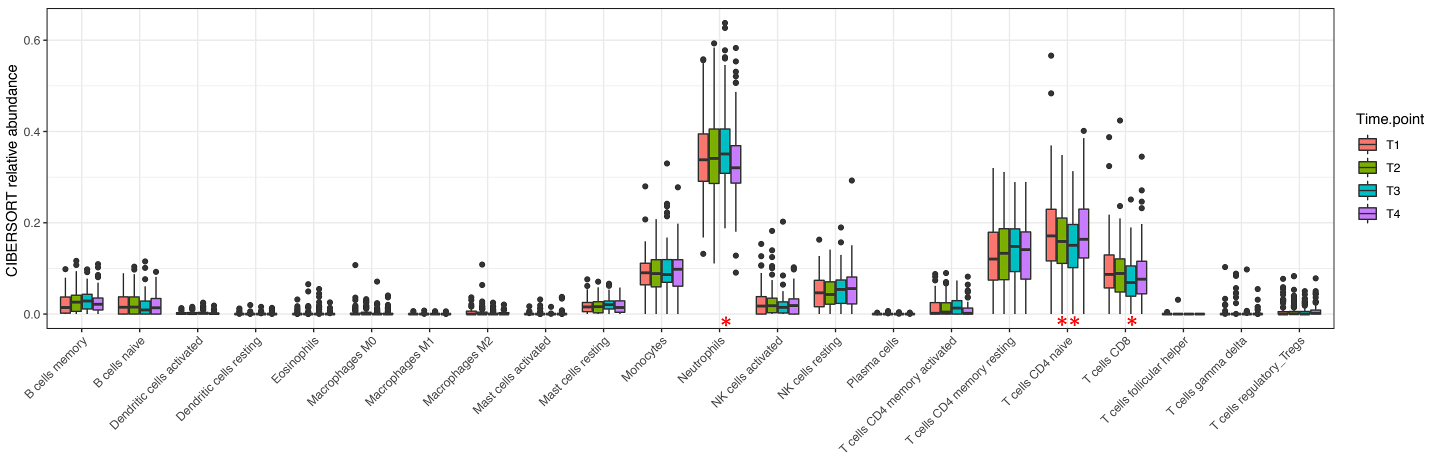

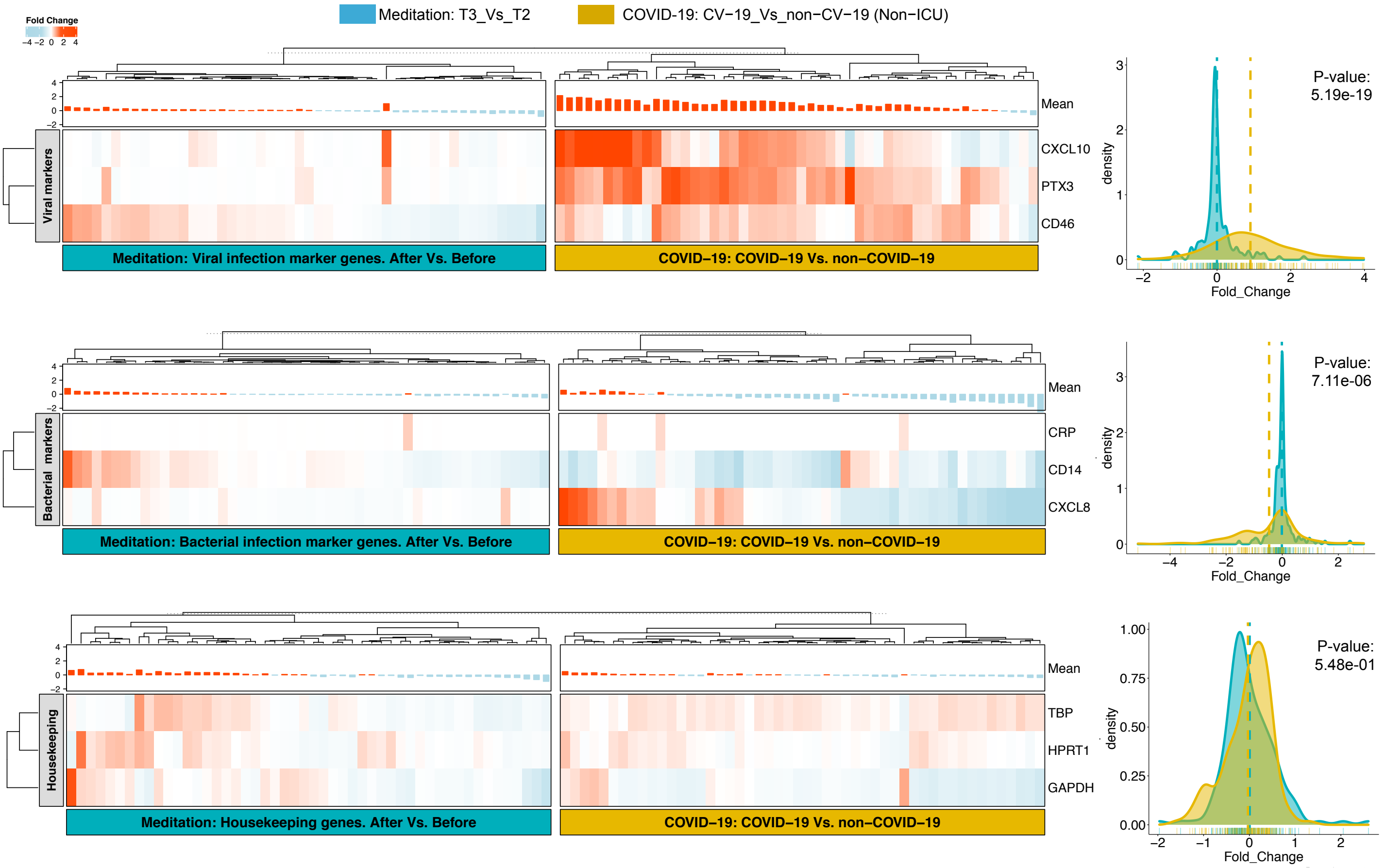

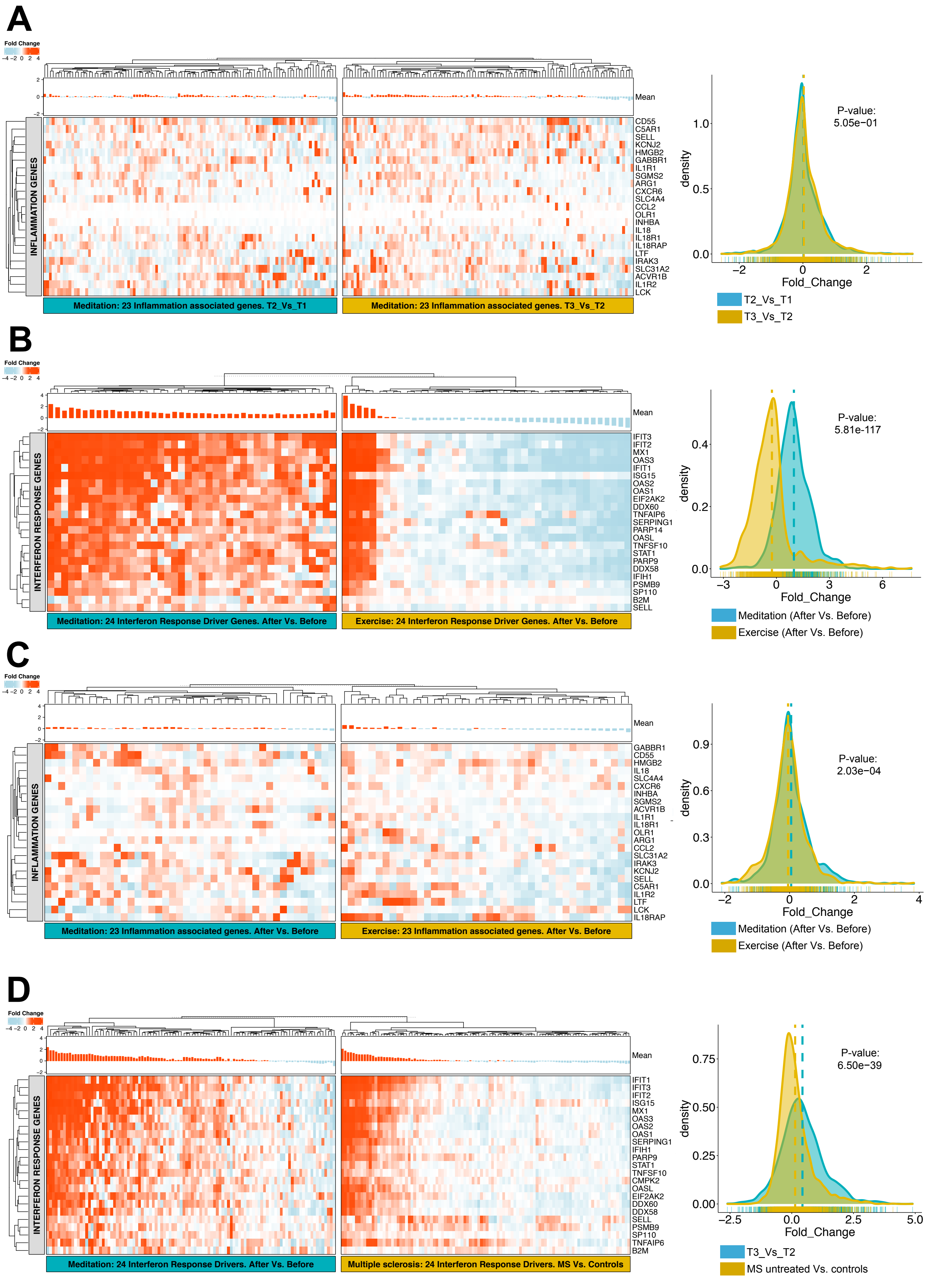
