## Supplemental_Figure_Legends for "Large-scale genomic study reveals robust activation of the immune system following advanced Inner Engineering meditation retreat"

SUPPLEMENTARY FIGURE/TABLE LEGENDS:

**Fig. S1 The quantity of 22 different immune cell types estimation in the meditation gene expression data using CIBERSORT-Relative deconvolution.** (A) The box plot visualizing the relative abundance of 22 immune cell subsets. CIBERSORT-derived immune cell relative scores were used to determine the abundance of immune cells in all the samples derived from four-time pre-and post-meditation (T1-T4). Significantly higher abundance of neutrophils and significantly lower abundance of CD8+ T-cells and naïve CD4+ T-cell was observed at T3 compared to T1. Comparisons were performed by employing two-way ANOVA test. *P < 0.05.

**Fig. S2 Comparison of meditation and COVID-19 transcriptional profiles showing viral and bacterial marker genes.** Heatmaps are depicting the expression of the three bona fide viral markers (top), bacterial markers (middle), and control genes (bottom) across meditation (left - T3 (after meditation) versus T2) and COVID-19 (right - non-ICU COVID-19 patients versus non-ICU non-COVID-19 patients) samples (columns). For all heatmaps, the red color corresponds to gene upregulation and blue to downregulation. Mean gene expression levels are shown as a bar-plot on top of each heatmap. All the heatmaps are supplemented with density plots showing the distribution of log2 fold change, and the significance of the variability in the expression levels between the two groups are calculated by a two-sample t-test.

**Fig. S3 Comparison of meditation, exercise, and multiple sclerosis specific transcriptional profiles in terms of interferon and inflammatory response.** (**A**) Heatmaps are depicting the expression of the 23 inflammatory response genes (rows) across meditation samples (columns) for two different timepoint comparisons, T2 (before meditation) versus T1 (baseline) and T3 (after meditation) versus T2, showing no significant changes. (**B**) Heatmaps are comparing the expression of the 24 meditation-specific IFN driver genes across meditation samples (left) and in samples obtained after exercise training (right) for two different comparisons; after meditation (T3) versus before meditation (T2) and after exercise training versus before exercise training. (**C**) Heatmaps comparing the expression of the 23 inflammatory response genes in samples before and after meditation and exercise training for two different comparisons as described in **B**. (**D**) Heatmaps are comparing the expression of the 24 meditation-specific IFN driver genes across meditation samples (left) and in samples obtained from multiple sclerosis patients (right) for two different comparisons; after meditation versus before meditation and MS patients versus controls. For all heatmaps, the red color corresponds to gene upregulation and blue to downregulation. Mean gene expression levels are shown as a bar-plot on top of each heatmap. All the heatmaps are supplemented with density plots showing the distribution of log2 fold change, and the significance of the variability in the expression levels between the two groups are calculated by a two-sample t-test.

**Table S1.** Differentially expressed genes before and after the meditation retreat at four-time points. Genes with a significant differential expression in batch and cell type composition corrected data in all four timepoints (T1, T2, T3, T4) by pair-wise comparison of all six permutation combinations are provided.

**Table S2.** Gene-module membership association based on WGCNA co-expression networks. Nine robust modules identified from datasets generated at four time points before and after meditation are denoted along with genes in these modules, the gene significance values (correlation of a gene expression profile with a sample trait) with T3, and the module membership values (Intramodular connectivity).

**Table S3.** Gene ontology enrichment analysis of meditation associated modules. For gene set enrichment analysis for GO terms, we considered GO terms with Fisher's Exact Test P-values less than 0.05. Enriched GO terms are provided for each module in separate tabs.

**Table S4.** NetBID analysis identifies meditation-associated drivers. Ninety drivers with significant differential activity in comparing T3 versus other time points are provided in this table, along with the differential gene expression values.

**Table S5.** Gene set enrichment analysis of meditation-associated drivers. Gene set enrichment analysis against the collection of annotated gene sets from MSigDB was utilized to elucidate the functional relevance of ninety meditation-associated drivers. We considered gene sets with Fisher's Exact Test P-values less than 0.05 as enriched. Enriched gene sets terms are provided for up and down drivers in separate tabs.

**Table S6.** Analysis of transcription-factor binding-sites (TFBS) enrichment in the meditation specific co-expression modules. For estimation of TFBSs enrichment in the identified corresponding module-genelist (top 200 genes based on connectivity) promoter sequences (1000bp upstream from transcription start site), P-values were obtained relative to three background datasets: 1000‐bp of sequence upstream of all human gene, human CpG islands and human chromosome 20 (see methods). Enriched TFBS position weight matrices from both JASPAR and HOCOMOCO databases are provided in this table. Enriched TFBSs are provided for each module in separate tabs.

**Table S7.** Literature annotation of enriched transcription factors associated with interferon signaling. Table providing enriched transcription factor associated with interferon signaling based on the published literature by testing association with the key-words: ‘interferon signaling’ and ‘interferon pathway’ in the PubMed database for every gene. The total number of hits (publications) for each gene is represented.
